# *Aedes aegypti* gut transcriptomes respond differently to microbiome transplants from field-caught or laboratory-reared mosquitoes

**DOI:** 10.1101/2023.03.16.532926

**Authors:** Shivanand Hegde, Laura E. Brettell, Shannon Quek, Kayvan Etebari, Miguel A. Saldaña, Sassan Asgari, Kerri L. Coon, Eva Heinz, Grant L. Hughes

## Abstract

The mosquito microbiome is critical for host development and plays a major role in many aspects of mosquito biology. While the microbiome is commonly dominated by a small number of genera, there is considerable variation in composition among mosquito species, life stages, and geography. How the host controls and is affected by this variation is unclear. Using microbiome transplant experiments, we asked whether there were differences in transcriptional responses when mosquitoes of different species were used as microbiome donors. We used microbiomes from four different donor species spanning the phylogenetic breadth of the Culicidae, collected either from the laboratory or field. We found that when recipients received a microbiome from a donor reared in the laboratory, the response was remarkably similar regardless of donor species. However, when the donor had been collected from the field, far more genes were differentially expressed. We also found that while the transplant procedure did have some effect on the host transcriptome, this is likely to have had a limited effect on mosquito fitness. Overall, our results highlight the possibility that variation in mosquito microbiome communities are associated with variability in host-microbiome interactions and further demonstrate the utility of the microbiome transplantation technique.

## Background

The collection of microorganisms associated with an organism (*i*.*e*., its microbiome) has profound effects on its host biology. The mosquito microbiome in particular is critical for larval development (Coon et al., 2014), plays a profound role in host fitness (Giraud et al., 2022; Schmidt and Engel, 2021; Sharma et al., 2013), and, importantly, can affect the mosquito’s ability to transmit pathogens such as dengue and Zika viruses (Cansado-Utrilla et al., 2021; Carlson et al., 2020; Ramirez et al., 2012). As such, manipulating the mosquito microbiome has the potential to reduce transmission of globally important mosquito-borne pathogens.

Traditionally, manipulating the microbiome has involved treating mosquitoes with antibiotics that alter microbiome composition, but can also affect mosquito physiology (Chabanol et al., 2020; Ha et al., 2021). However, approaches rearing axenic (germ-free) mosquito larvae followed by supplementation with bacteria of choice have proven to be an excellent way to interrogate host-microbe interactions without using antibiotics, thus removing effects of the antibiotic and the ‘original’ microbiome. Largely, this gnotobiotic approach has been used for investigating the role of the microbiome in mosquito development (Coon et al., 2016; Correa et al., 2018). More recently, this approach has been exploited to perform interspecies microbiome transfers opening up the possibility to study microbial symbiosis in mosquitoes (Coon et al., 2022; Romoli et al., 2021).

The ability to rear axenic/gnotobiotic mosquitoes also provides an opportunity to understand how the presence or absence of gut microbial communities affect host gene expression. Previously, in a comparison of axenic, gnotobiotic and conventionally-reared *Aedes aegypti*, 1328 host transcripts were differentially expressed compared to gnotobiotic and conventionally-reared mosquito larvae (Vogel et al., 2017). However, a different study found a much smaller effect in adult *Ae. aegypti*, with only 170 genes differentially expressed between axenic and conventionally-reared mosquitoes (Hyde et al., 2020). These studies demonstrate the utility of the axenic/gnotobiotic system for investigating mosquito-microbiome interactions, and furthermore point to larval stages being key for understanding how the host reacts to the microbiome.

Recently, we developed an interspecies microbiome transplantation technique in mosquitoes and showed that we could successfully recapitulate microbial composition in the recipient host (Coon et al., 2022). This novel approach allowed us to manipulate the microbiome and to investigate the impact of complex heterogeneous communities on mosquito gene expression. This study sought to address two questions: (1) How does the *Ae. aegypti* transcriptome change upon receiving microbiome transplant when a different mosquito species is used as a microbiome donor? and (2) Does *Ae. aegypti* experience transcriptomic changes associated with the transplant procedure itself? To address the first question, we performed inter-species microbiome transplants using microbiomes from three donor species (*Aedes taeniorhynchus, Culex tarsalis* and *Anopheles gambiae*) and performed RNA-Seq analysis to compare recipient host transcriptional profiles to *Ae. aegypti* recipients transplanted with their original microbiome. We also considered whether microbiomes derived from field-caught or laboratory-reared *Ae. aegypti* and *Ae. taeniorhynchus* mosquitoes affect recipient host transcriptomes differently. To address the second question, we compared transcriptional profiles of each of the *Ae. aegypti* treatment groups that had received a microbiome transplantation to mosquitoes conventionally reared in the same system without a microbiome transplant. Using mosquito microbiome transplants to unravel the intricacies of how mosquitoes are affected by their microbiomes is relevant for both mosquito biology and our understanding of host-microbiome interactions more broadly.

## Methods

### Experimental setup

The experimental setup comprised seven treatments, each with three replicates (Figure 1): (*i*) *Ae. aegypti* receiving a transplant isolated from conspecific individuals of the same laboratory-maintained Galveston line (*i*.*e*., their original microbiome); *Ae. aegypti* receiving a transplant from one of five different donor pools from varying locations and phylogenetically distinct species (henceforth termed ‘extraneous donors’); these included (*ii*) field-caught *Ae. aegypti*, (*iii*) field-caught *Ae. taeniorhynchus*, (*iv*) laboratory-reared *Ae. taeniorhynchus*, (*v*) laboratory-reared *Cx. tarsalis*, and (*vi*) laboratory-reared *An. gambiae*; and (*vii*) *Ae. aegypti* Galveston line reared under aseptic conditions without egg sterilization to retain their original microbiome (conventionally-reared control).

**Figure 1.**
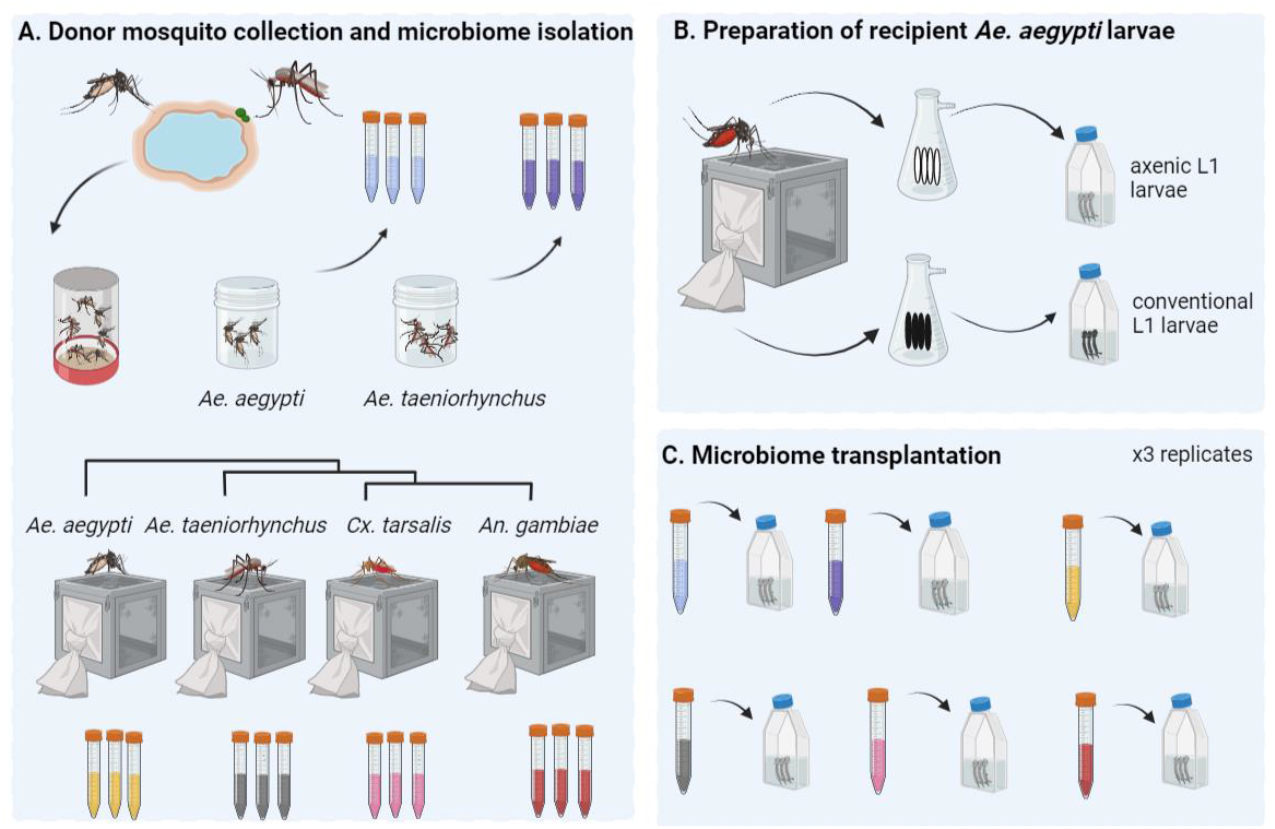
Microbiome transplantation from field-collected and laboratory-reared mosquitoes into recipient laboratory-reared mosquitoes. **A**. Adult mosquitoes from field populations of *Ae. aegypti* or *Ae. taeniorhynchus* were trapped using BG sentinel traps in Galveston, Texas and sorted according to species and sex. Three replicate pools of 20 adult females were then used to isolate donor microbiomes from each species. Donor microbiomes were also isolated from three replicate pools of 20 laboratory-reared *Ae. aegypti, Ae. taeniorhynchus, Cx. tarsalis*, and *An. gambiae* adult females. **B**. Laboratory-reared *Ae. aegypti* were used as recipient hosts for all transplants. In brief, eggs were surface sterilized using ethanol and bleach before vacuum hatching to obtain L1 axenic larvae. As a control for the transplantation process, we also vacuum hatched a batch of non-sterilized eggs from the same colony. These were grown conventionally in closed conditions to retain their original microbiome. (**C**) Axenic larvae were transferred into T75 tissue culture flasks at 20 larvae per flask with three replicates per treatment. Here they were inoculated with the donor microbiome through supplementation of the larval water. Flasks were maintained at 28 °C and fed with sterile fish food on alternative days. Once larvae had reached the fourth instar they were harvested, their guts dissected and RNA-Seq was carried out using pools of five guts for each of three replicate flasks per treatment. Figure created using Biorender.

### Donor mosquito collections

Microbiome transplantations were carried out by first isolating donor microbiomes from one of four mosquito species (*Ae. aegypti, Ae. taeniorhynchus, Cx. tarsalis*, or *An. gambiae*), which had either been laboratory-reared or field-caught (Figure 1). Colonies of all four species had been continually maintained at the University of Texas Medical Branch at 28 °C with 12 hr light/dark cycles and provided 20% sugar solution *ad libitum*. The laboratory colony of *Ae. aegypti* (Galveston line) were the F3 generation, whereas all other laboratory-reared mosquito colonies had been maintained for approximately ten years. Pools of 20 three-to-four-day old sugar fed adult females from one colony of each species were used for microbiome isolations. We also collected members of two of these species, *Ae. aegypti* and *Ae. taeniorhynchus* from field populations. Collections were made in 2018 locally in Galveston, Texas using Biogents sentinel (BG) traps. Adult mosquitoes were collected and sorted morphologically according to species and sex. Again, pools of 20 females of each of the two species were used for microbiome isolations.

### Preparation of recipient mosquitoes and microbiome transplantation

Microbiome isolation and transplantation was carried out using our recently developed methodology (Coon et al., 2022) as follows: Recipient mosquitoes were prepared by surface sterilising *Ae. aegypti* eggs using 70% ethanol and vacuum hatching under sterile conditions to generate axenic first instar larvae. The larvae were then transferred to T75 tissue culture flasks in sterile water at the rate of 20 larvae per flask (three replicate flasks per treatment). The same laboratory-reared *Ae. aegypti* (Galveston line) colony as used for microbiome donation was used as the source of recipient hosts for all transplants. For each of the six donor types (four laboratory-reared and two field-caught), three replicate pools of 20 mosquitoes were surface sterilised using 70% ethanol and bleach washes followed by homogenisation and filtration. Resulting donor microbiome aliquots were transplanted into recipient larvae by inoculating the larval water, with one aliquot per replicate flask. Recipient larvae were maintained in a closed environment at 28 °C with 12 hr light and dark cycle and supplemented with sterile fish food on alternative days until they reached the fourth instar. Since *Ae. aegypti* larvae require bacteria for their development (Coon et al., 2014), only those individuals that had been successfully inoculated with the donor microbiota developed.

### Sample preparation, RNA extraction and preparation of cDNA libraries for RNA-Seq

When recipient mosquitoes reached their fourth instar, five larvae were collected from each flask, surface sterilised, and their guts dissected. The five guts were then pooled to obtain sufficient RNA for cDNA library preparation and RNA-Seq. RNA was extracted using the PureLink RNA mini kit (Thermo Fisher Scientific), then using between 100ng-1ug total RNA, polyA+ RNA transcripts were isolated using the NEBNext Poly(A) mRNA Magnetic Isolation Module (New England Biolabs). Non-directional libraries were created using the NEBNext Ultra II RNA Library Prep Kit (New England biolabs) and Next Generation Sequencing was carried out using the Illumina NextSeq 550 platform to generate 75bp paired end reads at the University of Texas Medical Branch Core Next Generation Sequencing Facility.

## Data analysis

Sequence data were obtained in fastq format and quality checked using FASTQC v0.11.5 (Andrews, 2017). All samples had an average phred score of > 30, with no adapter sequences present so no trimming was performed. FeatureCounts v2.0.1 (Liao et al., 2014) was used to obtain raw count data from the sequencing files using default parameters and the *Ae. aegypti* reference genome (Genome version GCA_002204515.1, Annotation version AaegL.5.3) to determine feature locations. The resulting feature count table was then imported into RStudio v1.4.1106 and filtered to remove any genes which did not have at least ten reads present in each replicate of at least one treatment group before continuing with subsequent analyses.

Firstly, we investigated how *Ae. aegypti* responded to receiving a microbiome transplant from an extraneous donor. We compared gene expression in each recipient that had received a microbiome from a donor belonging to a different species, or from a different location to a baseline of recipients that had received a transplant of their ‘original’ microbiome from a conspecific donor. To focus on the gene expression in transplant-recipients, for this analysis we had removed the conventionally reared control mosquitoes. Differential expression (DE) analysis was carried out using DESeq2 v1.30.1 (Love et al., 2014) using default parameters. DESeq2 takes as input raw read counts from programs such as FeatureCounts, using the DESeqDataSetFromMatrix command. As part of its internal workflow, DESeq2 automatically normalizes gene expression data based on the input raw count data. Thresholds were applied to the resulting list of differentially expressed genes (DEGs) to retain only those with an adjusted p value of < 0.05 and an absolute log_2_ fold change of ≥ 1.5. An upset plot was created using the UpsetR package v1.4.0 (Conway et al., 2017) to visualise the number of DEGs in each pairwise comparison between recipients of a transplant from an extraneous donor and the recipients of a transplant from a conspecific donor, as well as show how many were common to multiple transplant groups or unique to one treatment. The ComplexHeatmap package v2.12.0 (Gu et al., 2016) was used to visualise the log_2_ fold changes of these DEGs compared to the ‘original’ microbiome control. We further investigated those DEGs identified as enhanced or suppressed when using each of the extraneous donor-derived microbiomes, by using the VectorBase Gene Ontology enrichment analysis tool to determine enriched GO terms (Biological Processes, Bonferroni adjusted p value < 0.05) in the enhanced or suppressed DEGs (VectorBase IDs).

To investigate how the recipient host transcriptome was affected by the transplant procedure itself, differential expression analysis was repeated using DESeq2 and conventionally reared mosquitoes as the baseline group to which all transplant groups were compared. The UpsetR and ComplexHeatmap packages were then used to compare DEGs present in every comparison to the conventional control and to plot associated log_2_ fold changes prior to GO enrichment analysis to identify functions of commonly enhanced and suppressed genes.

Sequencing reads were deposited in the National Centre for Biotechnology Information Sequence Read Archive under the accession PRJNA941184. All R code used in analyses, as well as raw counts table and metadata are available at https://github.com/laura-brettell/microbiome_transplant_RNASeq

## Results and Discussion

### Host gene expression shows marked differences when the microbiome donor was field-caught compared to laboratory-reared

Microbiome transplantation experiments provide a unique opportunity to investigate how the host interacts with a selection of diverse microbiomes in a controlled environment. Here, we used our previously developed methodology (Coon et al., 2022) to ask whether different microbiomes alter the host transcriptome. While mosquito microbiomes are commonly dominated by a small number of bacterial genera (Coon et al., 2014), microbiome composition varies amongst host species (Hegde et al., 2018; Kozlova et al., 2021), geography (Coon et al., 2016; Zouache et al., 2011), and across individuals (Coon et al., 2022; Osei-Poku et al., 2012). In our previous study, we found variability in the microbiome of three different mosquito species reared under identical insectary conditions (Hegde et al 2018). Hence, this begs question how do mosquitoes respond to these varied microbiomes.

To assess whether mosquitoes respond differently to varied mosquito-derived microbiomes, we performed transplantations using donors spanning the phylogenetic breadth of the Culicidae and a combination of laboratory-reared and field-caught samples. All microbiomes were transplanted into laboratory-reared *Ae. aegypti* (Galveston line) from the same generation (Figure 1). Larvae in all experimental treatments successfully developed to the fourth instar, indicating that each of the mosquito microbiomes used in this experiment provided the necessary nourishment for larval development. This is irrespective of donor species or collection environment, and is in agreement with the findings of several previous studies that looked at the impact of altered larval microbiomes on mosquito development (Correa et al., 2018; Vogel et al., 2017).

Using RNA-Seq, we compared gene expression in the guts of mosquitoes that received a microbiome from an extraneous donor (*i*.*e*., isolated from a different species or collected from a different environment) to those that received their original microbiome (*i*.*e*., isolated from conspecifics from the same *Ae. aegypti* laboratory population) (Figure 1). Across the entire dataset, we obtained an average of 23.6M reads per sample (range 16.1M – 30.8M) with an average of 74% of reads (range: 70.4% – 76.3%) mapping uniquely to the *Ae. aegypti* genome (Supplementary Table 1). Differential expression (DE) analysis revealed a striking difference between recipients of inoculated with laboratory-reared versus field-caught donor microbiomes. When recipients received a transplant from a donor reared in the same laboratory, there was little change to the gut transcriptome regardless of which donor species was used (Figure 2). Transplants using microbiomes derived from laboratory-reared *Ae. taeniorhynchus, Cx. tarsalis*, and *An. gambiae* donors resulted in 55, 49, and 19 DEGs, respectively (Figure 2, Supplementary Table 2). In contrast, transplantation using microbiomes derived from field-caught donors resulted in far more modulated transcripts, with microbiomes from field-caught *Ae. aegypti* resulting in 447 DEGs and those from field-caught *Ae. taeniorhynchus* resulting in 448 DEGs.

**Figure 2.**
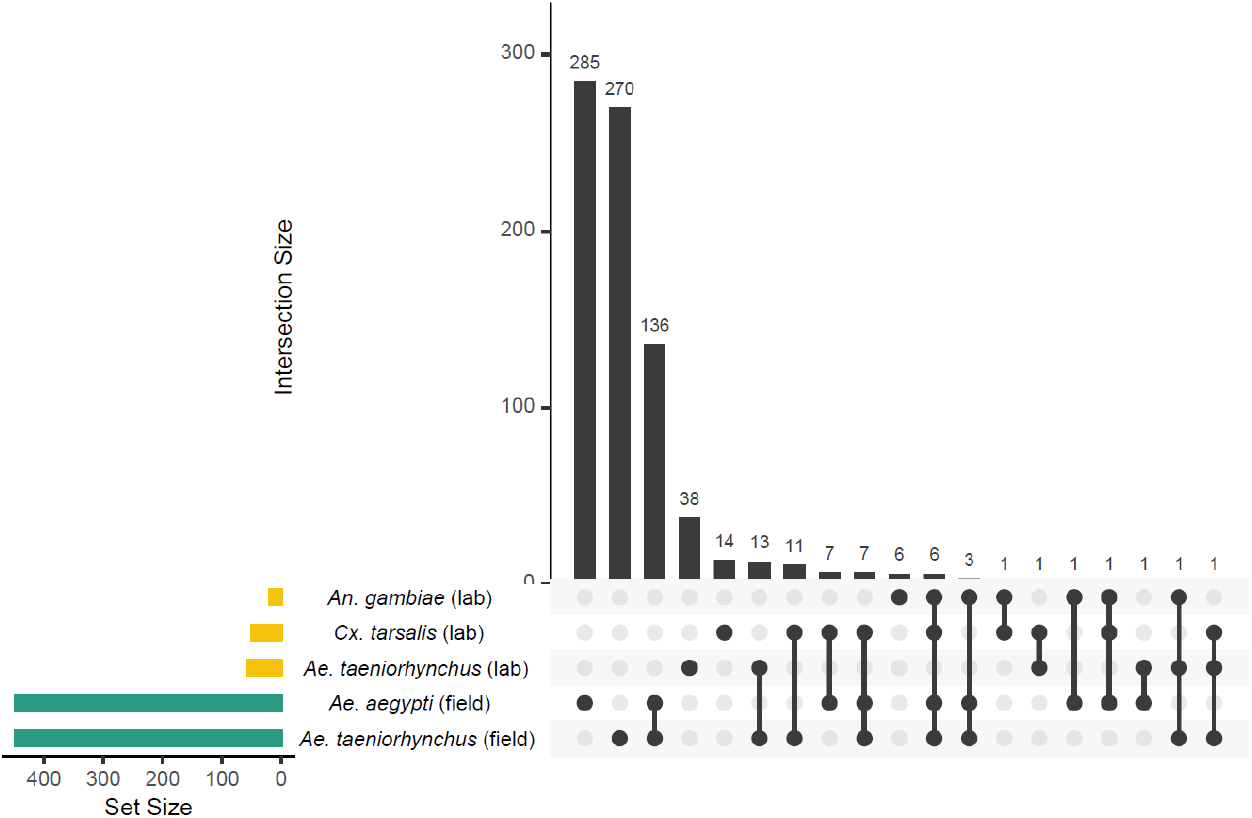
Upset plot showing the number of differentially expressed genes (DEGs) in each of the microbiome transplant recipients relative to the control recipients, that had received their original microbiome. Set size refers to the number of DEGs in the recipient when transplanted with microbiomes from each of five donor types (*An. gambiae, Cx. tarsalis*, and *Ae. taeniorhynchus* reared in the laboratory (yellow bars); and *Ae. aegypti* and *Ae. taeniorhynchus* collected from the field, (green bars)). Intersections where DEGs were identified in multiple transplantation types are denoted by the ball and stick diagram, with black bars showing the number of DEGs in each intersection, *i*.*e*., 285 DEGs were seen only when *Ae. aegypti* (field)-derived microbiomes were used.

While we did not characterize the composition of the different donor microbiomes in our study, the consistency in response, or lack thereof, of recipient hosts to laboratory-reared donor microbiomes suggests some level of similarity in composition between the different laboratory-derived donor microbiomes we isolated. The overall stronger differences in responses we observed across recipients of field-caught donor microbiomes also suggests that field-caught mosquitoes harbour more variable microbial communities that differ in composition from those present in laboratory-reared mosquitoes. This is also consistent with previous studies comparing the microbiomes of *Ae. aegypti* and other animals maintained in captivity to their free-living counterparts (Eichmiller et al., 2016; Lemieux-Labonté et al., 2016). Collectively, this suggests that microbiome composition is generally affected more by environment than host species, although it is not always the case (Hegde et al., 2018), which indicates that the factors governing microbiome assembly are complex. In each of the groups receiving a transplant from a field-caught donor, approximately one quarter of DEGs compared to the original microbiome control were common to both comparisons (136/447 for *Ae. aegypti* field donor and 136/448 for *Ae. taeniorhynchus* field donor) (Figure 2). We assume that the two field-derived microbiomes were different from one another, given we have previously seen that different species harbour distinct microbiomes (Hegde et al., 2018). However, the overlap in DEGs suggests some level of commonality in response, or that divergent field bacterial elicit similar transcriptional effects. Furthermore, of the DEGs common to both field-derived transplants, all but one DEGs showed the same direction of change (Supplementary Figure 1, Supplementary Table 2). Nine genes were enhanced when a transplantation was performed using a field-caught donor: a putative cytochrome b5 gene (AAEL004450), a ubiquitin-conjugating enzyme (AAEL001208), transcription initiation factor RRN3 (AAEL012265), a sterol o-acyltransferase (AAEL009596), and five for which the product is unknown. The same sterol o-acyltransferase has previously been found to be enhanced in gnotobiotic and axenically reared larvae compared to conventionally reared individuals (Vogel et al., 2017). Of the 126 genes that were suppressed in both field-transplant groups, 62 are of unknown function. However, the genes showing the strongest levels of suppression across the two field-transplant samples included three metalloproteases (AAEL011540 and AAEL011559, and the zinc metalloprotease AAEL008162). Zinc metalloproteases have previously been implicated as contributors to gut microbiome homeostasis in mice (Rodrigues et al., 2012). We did not identify any immune signal associated with receiving a microbiome transplant from an extraneous donor. Therefore, while immune function is affected by particular gut functions *i*.*e*., blood meal digestion (Hyde et al., 2020), it does not appear to be affected by the presence of different transplanted mosquito-derived microbiomes.

It is notable that when field-caught *Ae. taeniorhynchus* was used as the microbiome donor, similar numbers of genes were enhanced or suppressed compared to the original microbiome control (Figure 3). However, when using field-caught *Ae. aegypti* as the microbiome donor, recipients showed far greater numbers of suppressed than enhanced genes compared to the original microbiome control (Figure 3). That we did not observe a more profound effect when using field-caught *Ae. taeniorhynchus* donor microbiomes over field-caught *Ae. aegypti* donor microbiomes may be related to the inherent variability of using pools of field-caught mosquitoes.

**Figure 3.**
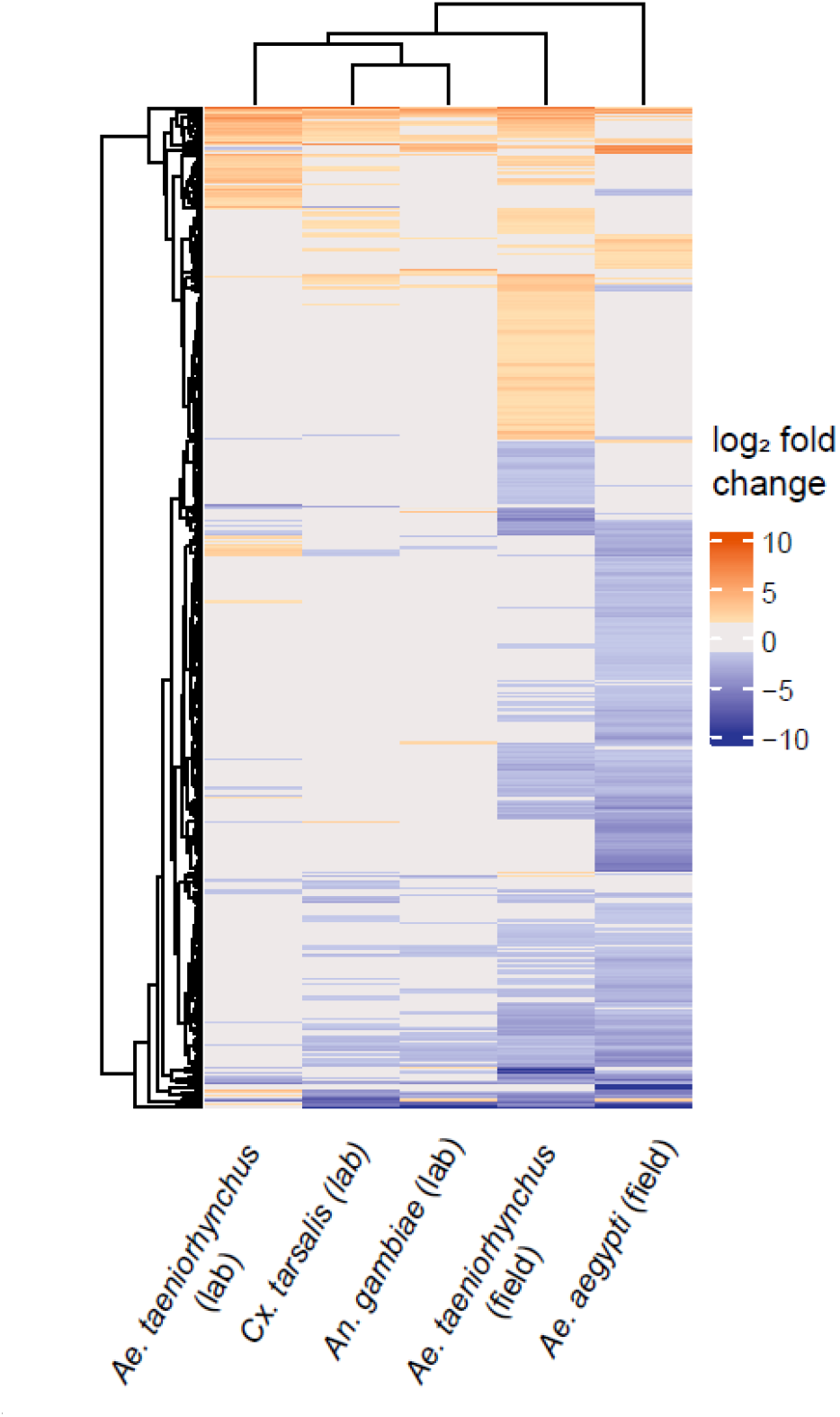
Heatmap showing differential gene expression between microbiome transplants using extraneous donors relative to transplants with laboratory-reared *Ae. aegypti* receiving their original microbiome. Orange cells represent when gene expression was enhanced in the transplant treatment (absolute log_2_ fold change ≥ 1.5, adjusted p value < 0.05). Blue cells represent a suppression of gene expression, passing the same thresholds. Grey denotes where a gene did not pass the differential expression threshold (log_2_ fold change > 1.5, adjusted p value < 0.05). The microbiome donor is shown on the *x*-axis, with each row on the *y*-axis corresponding to a DEG. The dendrograms represent clustering of similar responses as determined through the *hclust* function within the *ComplexHeatmap* package.

Given that the majority of DEGs were different between the two field-caught microbiome donor groups, we also looked at each of the two groups separately to identify whether any of the same biological processes may be implicated across both groups. We used Gene Ontology Enrichment Analysis to identify GO terms that were enriched in enhanced or suppressed DEGs in recipients of each of the field-derived microbiomes. Four biological processes were identified as suppressed in the recipients of both the *Ae. aegypti* (field) and *Ae. taeniorhynchus* (field) microbiomes (Supplementary Table 3). These include carbohydrate metabolic process, a dominant process of the anterior midgut and proventriculus (Hixson et al., 2022), transmembrane transport, obsolete oxidation-reduction process, and small molecule catabolic process. In keeping with the gene-level results, which showed only a small number of enhanced genes in the recipients of field-caught *Ae. aegypti* donor microbiomes, no GO terms were significantly enhanced. The recipients of field-caught *Ae. taeniorhynchus* donor microbiomes however, showed an enhancement of GO terms related to translation, including ribosome biogenesis, rRNA processing, and rRNA metabolic process.

### A core set of genes were consistently affected when conducting a microbiome transplantation

To maximise the potential of microbiome transplantation experiments, it is important to determine whether the transplant technique itself may influence the host. We know that transplant recipients successfully develop to adulthood (Coon et al., 2022), but we do not know if the recipients experienced transcriptomic changes associated with the experimental procedure. To address this, we compared the gut transcriptomes of *Ae. aegypti* larvae receiving a microbiome transplant (either their original microbiome or from a ‘foreign’ donor) to the gut transcriptomes of *Ae. aegypti* larvae from the same laboratory population that had not received a transplant to look for commonalities between responses (Figure 1).

We conducted differential expression analysis to compare gene expression in the conventionally reared larvae and each of the microbiome transplant treatments individually. We found 1680 DEGs in at least one transplantation group relative to the conventional control (Figure 4, Supplementary Table 4). This number ranged from 614 DEGs in the comparison between conventionally reared larvae and recipients of a field-caught *Ae. taeniorhynchus* donor microbiome, and up to 1269 genes in the comparison with recipients of a laboratory-reared *Ae. taeniorhynchus* donor microbiome. We then identified 71 genes that were consistently differentially expressed during each microbiome transplant, and thus could be a conserved response to the technique itself. Interestingly, these genes all showed the same direction of change in all comparisons, with 50 genes consistently enhanced when a transplant was performed, and 21 genes consistently suppressed (Supplementary Figure 2, Supplementary Table 5). Of the DEGs that were enhanced in the transplant recipients, one gene showed substantially higher differential expression than any other, a threonine dehydratase/deaminase gene (AAEL003564) involved in ammonia transport and detoxification (Durant et al., 2021). Among the most strongly suppressed DEGs in the transplantation groups were two glucosyl/glucuronosyl transferases (AAEL008560 and AAEL010381), genes previously found to be enriched in the L3/L4 life stages (Matthews et al., 2018).

**Figure 4.**
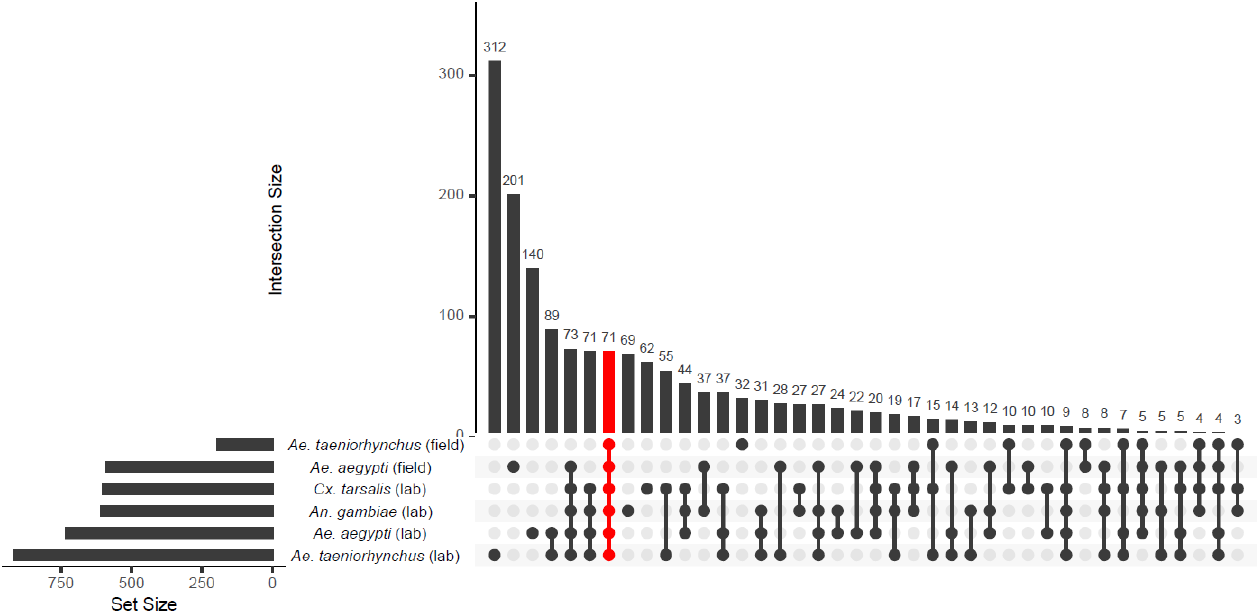
Upset plot showing the number of differentially expressed genes (DEGs) in recipients of each of the microbiome transplant treatments relative to the conventionally reared control. The 71 DEGs identified in every transplantation group are highlighted in red.

Given that the 71 genes identified in every comparison with conventionally reared controls were consistently affected in the same manner, we next asked whether other genes that had been identified in multiple comparisons were also affected in the same direction. We looked at all genes that passed our differential expression thresholds for at least one comparison and saw that, of the 1680 genes, all but 26 genes showed the same direction of change when they were identified in multiple comparisons (Figure 5, Supplementary Table 4). Thus, while only a small number of genes were identified in every comparison (and are therefore likely those most impacted by the transplant technique itself), there were general similarities in transcriptomic responses to a transplant overall. However, the magnitude of DEG changes between transplant recipients and conventionally reared controls varied amongst treatment groups. Interestingly, the treatment that showed the most similar transcriptome to conventional was the transplant using donor microbiomes isolated from field-caught *Ae. taeniorhynchus*, which as a different mosquito species and collection environment presumably harboured a substantially different microbiome composition to the *Ae. aegypti* control mosquitoes that were conventionally reared in the laboratory.

**Figure 5.**
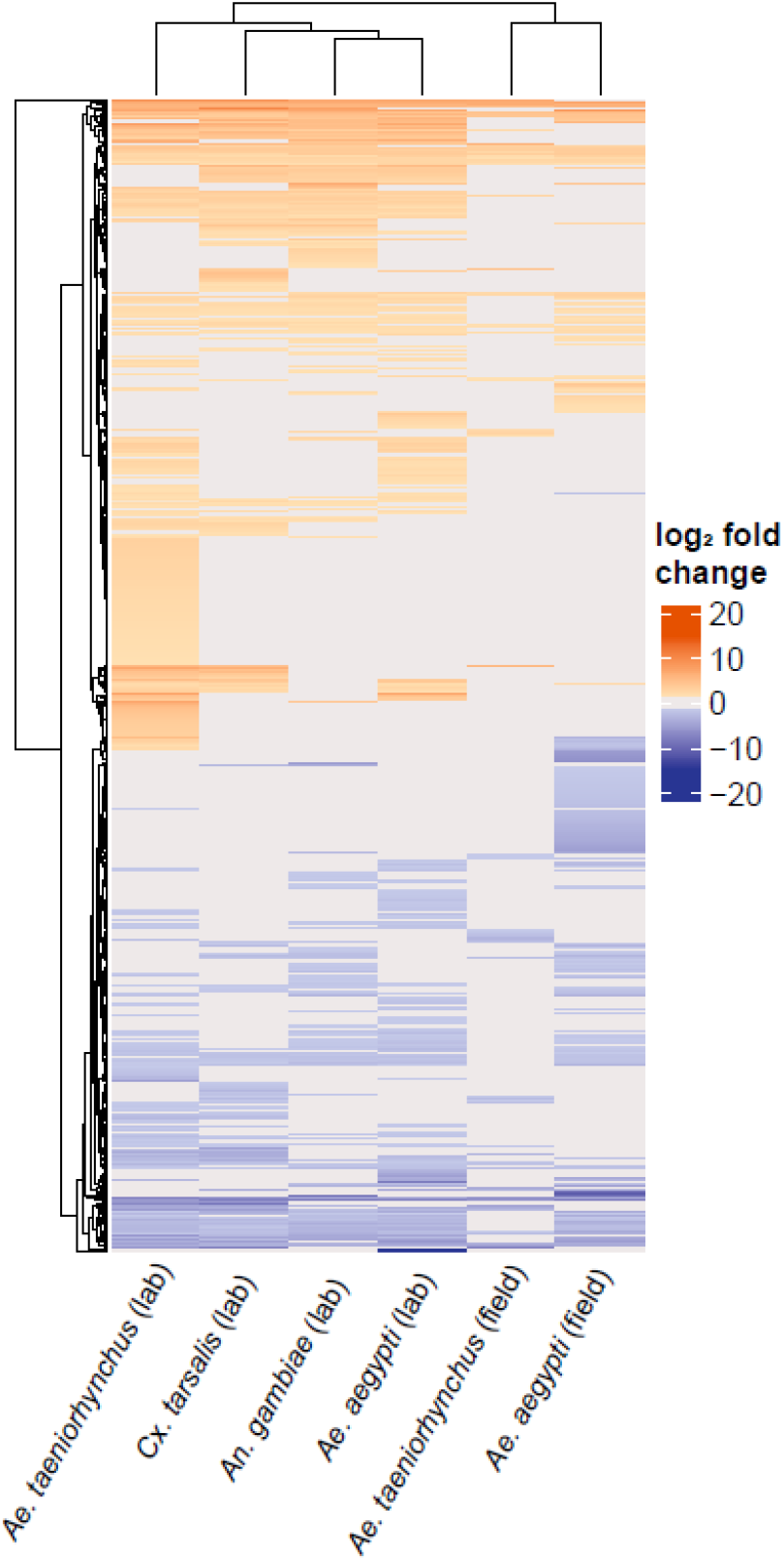
Heatmap showing the log_2_ fold change of each of the 1680 genes identified as differentially expressed in at least one comparison between a transplant treatment group and conventional. Warmer colours indicate when gene expression was enhanced in the transplant group and cooler colours indicate when gene expression was suppressed. Grey denotes where a gene did not pass the differential expression threshold (log_2_ fold change > 1.5, adjusted p value < 0.05). The microbiome donor is shown on the *x*-axis, with each row on the *y*-axis corresponding to a DEG.

To investigate whether biological functions could be implicated as being affected by the transplant process, we assigned GO terms to the genes that were consistently enhanced or suppressed in at least one transplant group across the dataset as a whole. The genes that were suppressed when a transplant was carried out were largely those with roles in metabolism and RNA processing (Supplementary Table 6), processes typically occurring in the gut (Hixson et al., 2022; Vogel et al., 2017). Indeed, one of the GO terms implicated in our data (ribonucleoprotein complex biogenesis) has previously been found to be affected by blood meal digestion (Hixson et al., 2022). Of the genes that were enhanced overall, when a transplant was performed, proteolysis was the only enriched GO term identified with a Bonferroni adjusted p value < 0.05.

Overall, these results support a lack of any strong, consistent physiological response to the transplant technique. While there were numerous DEGs identified amongst all different transplant groups compared to conventionally reared controls, most of these genes were only identified in a subset of comparisons. While other studies have shown alterations to the transcriptome when carrying out microbiome manipulations, there does not appear to be a consistent pattern. Hyde et al (2020) reported minimal effects on gut transcriptomes when comparing adult *Ae. aegypti* that had either received their native microbiome or been reared axenically. In contrast, Vogel et al (2017) reported a larger difference in the gut transcriptomes of first instar larvae that had been axenically or gnotobiotically reared compared to conventionally reared larvae. It should be noted that in both studies, these differences were likely attributable in large part to starvation stress associated with the developmental arrest of axenic larvae and are therefore not directly comparable to other studies, including this one, which sampled later life stages. Overall, we can speculate that while the transplant technique is likely having some effect, it is largely transient and not severely detrimental to the recipient host. Nevertheless, it is known that what bacteria mosquito larvae are exposed to can affect biological traits in adulthood (Carlson et al., 2020; Dickson et al., 2017), warranting further work to identify whether recipients are affected by the transplant technique as they develop into adulthood. Additionally, given our microbiome donors were all non-blood fed adults, it would be interesting to test what effect using donor microbiomes derived from this life stage had compared to other stages, including donor microbiomes derived from larvae or blood fed adults.

## Conclusions

The gut transcriptomes of *Ae. aegypti* responded differently to a microbiome transplant from a field-caught compared to a laboratory-reared donor, regardless of donor species. When the donor was laboratory-reared, even microbiomes derived from the most phylogenetically distant host showed a small number of DEGs. The responses imparted when a field-caught microbiome donor was used were far greater (more DEGs) and varied by donor species. The responses experienced across the transplants were varied and DEGs were generally those involved in normal gut functions such as metabolism. While we hypothesise that the responses seen here are not severely detrimental to the recipient mosquito, it does highlight the clear differences in microbiomes of laboratory-reared and field-caught mosquitoes, which must be considered when carrying out experiments with laboratory-reared mosquitoes. Taken together, these findings demonstrate the utility of the mosquito microbiome transplantation technique in determining the molecular basis of mosquito-microbiome interactions and underscores how mosquito larval life history has generally relaxed the dependence of larvae on any particular microbiome, at least under ideal diet/nutrient conditions. Future studies should focus on studying such interactions under variable diet/nutrient conditions that mimic field conditions and determining effects on adults.

## Acknowledgements

This work was supported by collaborative awards from the National Science Foundation and Biotechnology and Biological Sciences Research Council (NSF/2019368; BB/V011278/1) (to KLC, EH, and GLH) and National Institutes of Health (R21AI138074) (to GLH and KLC). KLC was further supported by the U.S. Department of Agriculture (2018-67012-29991). SH and LEB were supported by the LSTM Director’s Catalyst Fund. MS was supported by the NIAID Emerging and Tropical Infectious Diseases Training Program (5T32AI7526-17, PI: Lynn Soong)

## Supplementary Information

**Supplementary Table S1:** Summary of RNA-Seq data obtained, showing total number of paired reads for each sample with the proportion mapping to the *Ae. aegypti* reference genome (GCA_002204515.1), both singly and with multiple matches and the proportion of unmapped reads.

**Supplementary Table S2:** All differentially expressed genes that were identified in recipients of a microbiome transplant from an extraneous donor relative to control larvae that received their ‘original’ microbiome (passing thresholds of padj < 0.05 and absolute log_2_fold change ≥ 1.5). VectorBase IDs are given alongside log_2_ fold change when using each of the extraneous donors.

**Supplementary Table S3:** GO terms identified as enriched in differentially expressed genes that were enhanced/suppressed in recipients of an extraneous donor-derived microbiome, relative to control larvae that received their ‘original’ microbiome.

**Supplementary Table S4:** All differentially expressed genes that were identified in recipients of a microbiome transplant relative to control larvae that were conventionally reared in the laboratory (passing thresholds of padj < 0.05 and absolute log_2_fold change ≥ 1.5). VectorBase IDs are given alongside log_2_ fold change when a microbiome transplant was performed with each donor.

**Supplementary Table S5:** Differentially expressed genes that were commonly identified across all transplant groups relative to the conventionally reared control larvae (passing thresholds of padj <0.05 and log2fold change >1.5). VectorBase IDs and gene names are given alongside log_2_ fold change when larvae received microbiome transplants from each donor group.

**Supplementary Table S6:** GO terms enhanced/suppressed in recipients of microbiome transplants relative to a baseline of conventionally reared, no transplant controls.

## References

Andrews, S., 2017. FastQC: a quality control tool for high throughput sequence data. 2010.

Cansado-Utrilla, C., Zhao, S.Y., McCall, P.J., Coon, K.L., Hughes, G.L., 2021. The microbiome and mosquito vectorial capacity: rich potential for discovery and translation. Microbiome 9, 1–11.

Carlson, J.S., Short, S.M., Angleró-Rodríguez, Y.I., Dimopoulos, G., 2020. Larval exposure to bacteria modulates arbovirus infection and immune gene expression in adult Aedes aegypti. Dev. Comp. Immunol. 104, 103540.

Chabanol, E., Behrends, V., Prévot, G., Christophides, G.K., Gendrin, M., 2020. Antibiotic treatment in Anopheles coluzzii affects carbon and nitrogen metabolism. Pathogens 9, 679.

Conway, J.R., Lex, A., Gehlenborg, N., 2017. UpSetR: an R package for the visualization of intersecting sets and their properties. Bioinformatics.

Coon, K.L., Brown, M.R., Strand, M.R., 2016. Mosquitoes host communities of bacteria that are essential for development but vary greatly between local habitats. Mol. Ecol. 25, 5806–5826.

Coon, K.L., Hegde, S., Hughes, G.L., 2022. Interspecies microbiome transplantation recapitulates microbial acquisition in mosquitoes. Microbiome 10, 58. https://doi.org/10.1186/s40168-022-01256-5

Coon, K.L., Vogel, K.J., Brown, M.R., Strand, M.R., 2014. Mosquitoes rely on their gut microbiota for development. Mol. Ecol. 23, 2727–2739. https://doi.org/10.1111/mec.12771

Correa, M.A., Matusovsky, B., Brackney, D.E., Steven, B., 2018. Generation of axenic Aedes aegypti demonstrate live bacteria are not required for mosquito development. Nat. Commun. 9, 1–10.

Dickson, L.B., Jiolle, D., Minard, G., Moltini-Conclois, I., Volant, S., Ghozlane, A., Bouchier, C., Ayala, D., Paupy, C., Moro, C.V., Lambrechts, L., 2017. Carryover effects of larval exposure to different environmental bacteria drive adult trait variation in a mosquito vector. Sci. Adv. 3, e1700585. https://doi.org/10.1126/sciadv.1700585

Durant, A.C., Guardian, E.G., Kolosov, D., Donini, A., 2021. The transcriptome of anal papillae of Aedes aegypti reveals their importance in xenobiotic detoxification and adds significant knowledge on ion, water and ammonia transport mechanisms. J. Insect Physiol. 132, 104269.

Eichmiller, J.J., Hamilton, M.J., Staley, C., Sadowsky, M.J., Sorensen, P.W., 2016. Environment shapes the fecal microbiome of invasive carp species. Microbiome 4, 1–13.

Giraud, É., Varet, H., Legendre, R., Sismeiro, O., Aubry, F., Dabo, S., Dickson, L.B., Valiente Moro, C., Lambrechts, L., 2022. Mosquito-bacteria interactions during larval development trigger metabolic changes with carry-over effects on adult fitness. Mol. Ecol. 31, 1444–1460.

Gu, Z., Eils, R., Schlesner, M., 2016. Complex heatmaps reveal patterns and correlations in multidimensional genomic data. Bioinformatics 32, 2847–2849.

Ha, Y., Jeong, S., Jang, C., Chang, K., Kim, H., Cho, S., Lee, H., 2021. The effects of antibiotics on the reproductive physiology targeting ovaries in the Asian tiger mosquito, Aedes albopictus. Entomol. Res. 51, 65–73.

Hegde, S., Khanipov, K., Albayrak, L., Golovko, G., Pimenova, M., Saldana, M.A., Rojas, M.M., Hornett, E.A., Motl, G.C., Fredregill, C.L., 2018. Microbiome interaction networks and community structure from laboratory-reared and field-collected Aedes aegypti, Aedes albopictus, and Culex quinquefasciatus mosquito vectors. Front. Microbiol. 9, 2160.

Hixson, B., Bing, X.-L., Yang, X., Bonfini, A., Nagy, P., Buchon, N., 2022. A transcriptomic atlas of Aedes aegypti reveals detailed functional organization of major body parts and gut regional specializations in sugar-fed and blood-fed adult females. Elife 11, e76132.

Hyde, J., Correa, M.A., Hughes, G.L., Steven, B., Brackney, D.E., 2020. Limited influence of the microbiome on the transcriptional profile of female Aedes aegypti mosquitoes. Sci. Rep. 10, 1–12.

Kozlova, E.V., Hegde, S., Roundy, C.M., Golovko, G., Saldaña, M.A., Hart, C.E., Anderson, E.R., Hornett, E.A., Khanipov, K., Popov, V.L., Pimenova, M., Zhou, Y., Fovanov, Y., Weaver, S.C., Routh, A.L., Heinz, E., Hughes, G.L., 2021. Microbial interactions in the mosquito gut determine Serratia colonization and blood-feeding propensity. ISME J. 15, 93–108. https://doi.org/10.1038/s41396-020-00763-3

Lemieux-Labonté, V., Tromas, N., Shapiro, B.J., Lapointe, F.-J., 2016. Environment and host species shape the skin microbiome of captive neotropical bats. PeerJ 4, e2430.

Liao, Y., Smyth, G.K., Shi, W., 2014. featureCounts: an efficient general purpose program for assigning sequence reads to genomic features. Bioinformatics 30, 923–930.

Love, M.I., Huber, W., Anders, S., 2014. Moderated estimation of fold change and dispersion for RNA-seq data with DESeq2. Genome Biol. 15, 1–21.

Matthews, B.J., Dudchenko, O., Kingan, S.B., Koren, S., Antoshechkin, I., Crawford, J.E., Glassford, W.J., Herre, M., Redmond, S.N., Rose, N.H., 2018. Improved reference genome of Aedes aegypti informs arbovirus vector control. Nature 563, 501–507.

Osei-Poku, J., Mbogo, C., Palmer, W., Jiggins, F., 2012. Deep sequencing reveals extensive variation in the gut microbiota of wild mosquitoes from K enya. Mol. Ecol. 21, 5138–5150.

Ramirez, J.L., Souza-Neto, J., Torres Cosme, R., Rovira, J., Ortiz, A., Pascale, J.M., Dimopoulos, G., 2012. Reciprocal tripartite interactions between the Aedes aegypti midgut microbiota, innate immune system and dengue virus influences vector competence. PLoS Negl. Trop. Dis. 6, e1561.

Rodrigues, D.M., Sousa, A.J., Hawley, S.P., Vong, L., Gareau, M.G., Kumar, S.A., Johnson-Henry, K.C., Sherman, P.M., 2012. Matrix metalloproteinase 9 contributes to gut microbe homeostasis in a model of infectious colitis. BMC Microbiol. 12, 1–13.

Romoli, O., Schönbeck, J.C., Hapfelmeier, S., Gendrin, M., 2021. Production of germ-free mosquitoes via transient colonisation allows stage-specific investigation of host– microbiota interactions. Nat. Commun. 12, 1–16.

Schmidt, K., Engel, P., 2021. Mechanisms underlying gut microbiota–host interactions in insects. J. Exp. Biol. 224, jeb207696. https://doi.org/10.1242/jeb.207696

Sharma, A., Dhayal, D., Singh, O., Adak, T., Bhatnagar, R.K., 2013. Gut microbes influence fitness and malaria transmission potential of Asian malaria vector Anopheles stephensi. Acta Trop. 128, 41–47.

Vogel, K.J., Valzania, L., Coon, K.L., Brown, M.R., Strand, M.R., 2017. Transcriptome sequencing reveals large-scale changes in axenic Aedes aegypti larvae. PLoS Negl. Trop. Dis. 11, e0005273.

Zouache, K., Raharimalala, F.N., Raquin, V., Tran-Van, V., Raveloson, L.H.R., Ravelonandro, P., Mavingui, P., 2011. Bacterial diversity of field-caught mosquitoes, Aedes albopictus and Aedes aegypti, from different geographic regions of Madagascar. FEMS Microbiol. Ecol. 75, 377–389.

